# Sorghum Association Panel Whole-Genome Sequencing Establishes Pivotal Resource for Dissecting Genomic Diversity

**DOI:** 10.1101/2021.12.22.473950

**Authors:** J. Lucas Boatwright, Sirjan Sapkota, Hongyu Jin, James C. Schnable, Zachary Brenton, Richard Boyles, Stephen Kresovich

**Affiliations:** Department of Plant and Environmental Sciences, Clemson University, Clemson, SC 29634, USA; Advanced Plant Technology, Clemson University, Clemson, South Carolina 29634, USA; Center for Plant Science Innovation and Department of Agronomy and Horticulture, University of Nebraska-Lincoln, Lincoln, NE 68588, USA; Carolina Seed Systems, Darlington, SC 29532, USA; Pee Dee Research and Education Center, Clemson University, Florence, SC 29506; Feed the Future Innovation Lab for Crop Improvement, Cornell University, Ithaca, NY 14850

## Abstract

Association mapping panels represent foundational resources for understanding the genetic basis of phenotypic diversity and serve to advance plant breeding by exploring genetic variation across diverse accessions with distinct histories of evolutionary divergence and local adaptation. We report the whole-genome sequencing (WGS) of 400 sorghum [*Sorghum bicolor* (L.) Moench] accessions from the Sorghum Association Panel (SAP) at an average coverage of 38X (25X-72X), enabling the development of a high-density genomic-marker set of 43,983,694 variants including SNPs (~ 38 million), indels (~ 5 million), and CNVs (~170,000). We observe slightly more deletions among indels and a much higher prevalence of deletions among copy number variants compared to insertions. This new marker set enabled the identification of several putatively novel genomic associations for plant height and tannin content, which were not identified when using previous lower-density marker sets. WGS identified and scored variants in 5 kb bins where available genotyping-by-sequencing (GBS) data captured no variants, with half of all bins in the genome falling into this category. The predictive ability of genomic best unbiased linear predictor (GBLUP) models was increased by an average of 30% by using WGS markers rather than GBS markers. We identified 18 selection peaks across subpopulations that formed due to evolutionary divergence during domestication, and we found six F_st_ peaks resulting from comparisons between converted lines and breeding lines within the SAP that were distinct from the peaks associated with historic selection. This population has been and continues to serve as a significant public resource for sorghum research and demonstrates the value of improving upon existing genomic resources.

**Author summary:** 

## Introduction

Sources of natural genetic variation are foundational to crop improvement as they can be used in the genetic dissection of pivotal traits and the development of breeding populations. Along with advances in modern statistics and technology, maintenance and expansion of genetic diversity within available germplasm is paramount to the advancement of crop improvement. Since the early development of association mapping populations in maize [1], these panels have served to propel plant breeding by exploring genetic variation across diverse accessions with distinct histories of evolutionary divergence and local adaptation. Now, two decades after that initial maize panel, all major cereal grains have association panels including barley [*Hordeum vulgare* L.] [2], maize [*Zea mays*, L.] [1], rice [*Oryza sativa* L.] [3], sorghum [*Sorghum bicolor* (L.) Moench] [4, 5, 6], and wheat [*Triticum aestivum* L.] [7, 8]. Association mapping panels leverage the existing natural variation – both genetic and phenotypic – of a population to resolve complex trait variation to the genomic features influencing the phenotypic variance. As such, the diversity of germplasm in an association panel is vital to increase our understanding of causal biological mechanisms and translate to crop improvement [9].

The sorghum association panel (SAP), the first sorghum diversity panel, is composed of temperately adapted breeding lines, as well as converted (photoperiod-insensitive) tropical accessions from the Sorghum Conversion Program (SCP) [10, 11]. The accessions in the SAP were selected to maximize the genetic and phenotypic diversity of the panel as well as capture accessions that are important for understanding the demographic history and historical breeding importance based on known resistances or tolerances to abiotic and biotic stresses [4]. Sorghum’s broad geographic distribution [12, 4] and carbon-partitioning regimes [13] have resulted in two classification systems that distinguish accessions based on variation by race and carbon partitioning, with race representing the predominant classification system in the SAP. Sorghum is classified into five botanical races: bicolor, caudatum, durra, guinea, and kafir, which are thought to have formed through multiple domestication and adaptation events across different clines [14, 5].

The SAP was originally genotyped using simple sequence repeat (SSR) markers [4] and later sequenced using restriction-site based genotyping-by-sequencing (GBS) to obtain low-coverage single-nucleotide polymorphism (SNP) data [5, 15]. However, as sequencing costs have continued to decline, large association panels such as the SAP can be sequenced using whole-genome sequencing (WGS) at a lower cost to generate reliable genomic variants at high density for applications in genetics and breeding [16]. These high-throughput sequencing data can enable computational analyses for genomics-assisted breeding with the help of diverse variant types including single-nucleotide polymorphisms (SNPs), insertions, deletions (indels), and larger structural variants (SV). These variant types are also valuable for understanding genetic diversity when combined into variant graphs, also known as pan-genomes [17, 18]. Increased variant density also permits the identification of causal variants as opposed to variants that simply lie in linkage disequilibrium (LD) with causal variants. The application of WGS to the SAP will increase the power and utility of the SAP, just as GBS improved upon SSR markers, and serve to expand upon the identified genetic diversity that facilitates genome-wide association mapping (GWAS) and genomic selection (GS) in sorghum [19].

A variant graph better represents the true diversity of variant information across a population and ameliorates issues associated with mapping bias inherent in traditional reference-based genomics [18, 20]. When a traditional reference genome is used, variants missing from the reference, such as those arising from recent duplications or deletions, cannot be identified by QTL mapping or GWAS [21]. Such limitations are particularly pervasive when studying diverse accessions, and while using a different reference genome can circumvent this issue, variant graphs are particularly well-suited to capture this information and significantly reduce mapping biases [21]. Additionally, tools such as the GATK [22, 23] permit joint calling of variants across samples in a population to increase the power to detect true variants, and when indels are present, the joint calling methods can assess variants through localized assembly from the read data to reduce the impact of read-mapping biases on variant discovery [23]. Together, these tools permit robust variant discovery along with development of a diverse pan-genomic reference for future studies in sorghum.

In this study, we report the development and use of high-density genomic variants including SNPs, indels, and copy number variants (CNVs) for population and translational genomic analysis using whole genome sequencing (WGS). The application of high-throughput genotyping and robust variant discovery for the highly diverse SAP provides the genomic resources necessary for acceleration of gene discovery, genomics-assisted breeding, and genetic engineering toward improved cultivar development and carbon-negative agriculture. We demonstrate the value of the WGS genomic resource over GBS markers through comparative advantages in identification of novel genomic associations and increased accuracy in genomic prediction for various traits.

## Materials and methods

### Plant material and datasets

A total of 400 accessions in the United States Sorghum Association Panel (SAP) [4] were obtained through the Agricultural Research Service-Germplasm Resources Information Network (ARS-GRIN) (http://www.ars-grin.gov) (S1 File). Seedlings were grown by sowing 3-5 seeds from each accession in a plastic pot in the Biosystems Research Complex greenhouse at Clemson University, Clemson, SC. Tissue was collected from two-week-old seedlings and lyophilized for three days in a LABCONCO FreeZone 4.5L −50 °C benchtop freeze dryer prior to DNA extraction. Phenotypic data for all the traits used in genome-wide association and prediction analyses were derived from previously published datasets [15, 24, 25, 26]. We accessed the publicly available GBS data for the SAP to conduct comparative analyses between our WGS data to GBS marker data [5, 15].

### Whole genome sequencing data production and processing

Whole-genome sequencing (WGS) data was generated by RAPiD Genomics, Gainesville, FL using DNA extracted from lyophilized leaf tissue. WGS libraries were paired-end sequenced at ~30x coverage using an Illumina NovaSeq sequencer resulting in 2×150-bp reads. WGS reads were cleaned using fastp [27] before aligning with BWA version 0.7.17 [28] to the BTx623 version 3.1.1 annotated reference genome [29] obtained from Phytozome [30]. Both SNP and indel variants were called using the Genome Analysis Toolkit (GATK) pipeline version 4.1.7.0 [22] following GATK best practices [31, 32]. Joint calling in the GATK was used to increase sensitivity for low-frequency variants, better distinguish between homozygous reference sites and sites with missing data, and to maximize SNP fidelity by allowing accurate error modeling [23]. Variants were subsequently quality filtered using QD < 2.0, InbreedingCoeff < 0.0, QUAL < 30.0, SOR > 3.0, FS > 60.0, MQ < 40.0, MQRankSum < −12.5, and ReadPosRankSum < −8.0. Beagle version 5.1 was used to impute missing genotype data for biallelic SNPs in the variant call format (VCF) file resulting from the GATK pipeline. SNP density plots were generated using R-CMplot version 3.6.0 (https://github.com/YinLiLin/R-CMplot) in the R programming language [33].

The inbreeding coefficient and nucleotide diversity were calculated using VCFtools version 0.1.16 [34]. Nucleotide diversity was estimated using a non-overlapping 1 Mb sliding window and plotted using Circos [35]. The effects of SNPs and indels were predicted using snpEff [36] and general variant statistics were collected using BCFtools [37]. The variant metrics and predictions were collected and plotted using MultiQC [38]. GBS and WGS variant effect counts were collected from the snpEff results and plotted using Excel. To compare GBS and WGS coverage, SNPs were counted by 5, 10, 15, 20, and 40 kb bin sizes using a custom Python script and plotted using ggplot2 [39]. The linkage disequilibrium (LD) decay plot was generated using PopLDdecay v3.40 [40] and custom R scripts [33]. HaploBlocker v1.6.06 [41] was used to identify subgroup-specific haplotype blocks where a haploblock is defined as a sequence of genetic markers that occurs at least five times within the population. Each accession is then checked to determine if they contain a similar sequence of markers, which served to screen the population in a group-wise, identity-by-descent manner [41]. The number of ancestral populations represented by the SAP was estimated using the R package adegenet v2.1.3 [42] where a Discriminant Analysis of Principal Components (DAPC) was performed for 1-12 clusters and Bayesian Information Criteria was used to identify the optimal number of clusters to describe the population. Subsequently, ADMIXTURE v1.3.0 was executed using the number of clusters estimated from DAPC as K to visualize the degree of admixture across the SAP [43, 44].

Numerous tools can detect CNVs from WGS data, but the complexity of plant data can complicate accurate variant calling, especially when many tools were designed with default settings suited for human data [45]. As such, we called CNVs using Hecaton v0.3.0 [45], which uses multiple CNV tools, including DELLY v0.8.5 [46], GRIDSS v2.0.1 [47], LUMPY v0.2.13 [48], and Manta v1.4.0 [49], to detect CNVs before using a random forest model to distinguish probable false-positive from true-positive variant calls based on a pretrained model specific to plants. Hecaton has been shown to outperform current methods when applied to short-read WGS data of Arabidopsis, maize, rice, and tomato [45].

### Genome-wide analysis for selection signatures

Subpopulations identified using ADMIXTURE analysis (K=6) were used to estimate F_st_ according to the methods of [50] using the vcftools function --weir-fst-pop on the SNP variants after filtering for minor allele frequency > 5% for each subpopulation [34]. A window size of 1 Mb with a step size of 100 kb was used for calculations. F_st_ estimates were calculated for each subpopulation against all other subpopulations, and mean F_st_ for a subpopulation at a genomic window was computed as averaged F_st_ of a subpopulation against all other subpopulations for that genomic window. Additionally, we computed F_st_ between accessions derived from the sorghum conversion program (N=240) and temperate breeding lines (N=96) within the panel using the same parameters mentioned above (S1 File). Tajima’s D for the whole panel was calculated for 1 Mb non-overlapping windows using the vcftools function *--TajD*.

### Genome-wide association studies

The software GEMMA v0.98.1 [51, 52] was used for GWAS. The VCF files containing biallelic SNPs, indels, or CNVs were converted to PLINK format using PLINK [53], and GEMMA was then used to calculate a standardized relatedness matrix for linear mixed modelling on the filtered data (--miss=0.3 --maf=0.05). All models were run using a MAF filter of 0.05, and LMMs of the following the form,

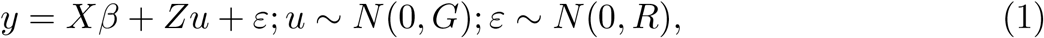

where y is a vector of phenotypic values for a single trait, X is a numeric genotype matrix generated from the variants, *β* represents an unknown vector of fixed effects and includes the effect size for each variant, Z is the design matrix for random effects, u is an unknown vector of random effects, and *∊* is the unknown vector of residuals. These models test the alternative hypothesis *H*_1_: *β* = 0 against the null hypothesis *H*_0_: *β* = 0 for each variant. Manhattan and Q-Q plots were generated using R-CMplot version 3.6.0 and ggplot2 version 3.3.5. GEMMA was also used to run Bayesian Sparse Linear Mixed Models (BSLMM) to better identify causative variants, with a probit model used for binary phenotypic data. The BSLMM model assumes fixed effects are distributed according to the sparse prior, 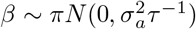[52]. Linkage disequilibrium (LD) statistics were calculated for significant loci using PLINK v1.9 [53].

A multivariate adaptive shrinkage approach was used to assess the degree of pleiotropic effects across the traits. Using the estimated effect sizes and standard errors for each marker from the GEMMA LMMs, a local false sign rate (lfsr) was calculated on a condition-by-condition basis using ashr in R [54] to filter variants based on a lfsr < 0.1. The lfsr represents the probability of incorrectly assigning the sign of an effect. The lfsr has been demonstrated to serve as a superior measure of significance over traditional multiple-testing corrections such as Bonferroni or False Discovery Rate [55] due to its general applicability and robust estimation process [54]. A control set of 1,200,000 random markers was also generated from the full set of markers to estimate the covariance between markers for each phenotype. From this control set, a correlation matrix was estimated using mashr [56] to control any confounding effects arising from correlated variation among the traits. Using both canonical and data-driven covariance matrices, we tested for pleiotropy across traits. Posterior probabilities were estimated for each marker using a mash model with all marker tests. The CDBNgenomics R package [57] was then used to extract Bayes factors and generate a Manhattan plot of the mash results where Bayes Factors > 3 were considered significant for pleiotropic effects.

### Genome-wide prediction

Previously published SAP phenotypic data [15, 24, 25, 26] were used to compare genomic prediction results between WGS and GBS marker data using the R package sommer [58]. A genomic best linear unbiased prediction (GBLUP) model of the following form was fit,

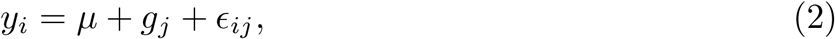

where *y_i_* is a vector of BLUPs for trait i, *μ* is the overall mean, *g_j_* is a vector of random effect of genotypes with 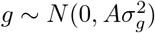, where 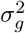 is the additive genetic variance and *A* is the realized additive relationship matrix calculated from a *n × m* genotype matrix with *n* genotypes and *m* markers using the *A.mat* function from the rrBLUP package [59], and *e_ij_* is a vector of residuals that are independent and identically distributed with 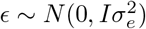, where 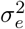 is the residual variance and *I* is an identity matrix.

Model performance was assessed using ten-fold cross validation, where individuals in nine of the folds were used for model training and the remaining fold was used as the testing set. Predictive ability was calculated as Pearson’s correlation coefficient between predicted and observed values for the testing set. A total of 100 iterations were run for each trait using the *set.seed* function in R for sampling seeds from 123 to 222. The predictive ability between GBS and WGS were compared using the pairwise t-test (*pairwise.t.test* function in R), and p-values were adjusted using Benjamini-Hochberg correction [55].

### Variant graph construction

A variant graph was constructed using *vg* [17], which incorporated the Sorghum BTx623 reference genome together with SPA variants (SNP, Indel, and CNV) called using the GATK [22] and Hecaton [45] pipelines. Individual chromosomes were constructed using vg construct with the options *--handle-sv* and *--node-max* of 32. After chromosome-level construction, each subgraph was given unique node identifiers using *vg ids* before building a single joint variant graph in XG format, which permitted querying of the variant graph and alignment of read data. Reads obtained from [60] were then aligned using the joint variant graph and *vg map* with default parameters. The resulting GAM file was quality filtered and used to calculate read support with *vg pack* before calling variants on individual samples with *vg call*.

## Results

### SNPs, indels, and copy number variants

We sequenced 400 accessions from the sorghum association panel (SAP) at an average of 38X coverage, ranging from a minimum of 25X up to 72X (S1 Fig, S2 Fig) and representing a total of approximately 82 billion reads or ~11 trillion bases after quality control. Reads exhibited highly consistent GC content across all samples (S3 Fig). The SAP genotypic data contained 43,983,694 variants, which includes SNPs (~ 38 million) and indels (~ 5 million) identified using GATK pipeline, and CNVs (~ 170,000) called using Hecaton (S1 Table). A total of 19,708,560 SNPs passed quality filtering based on variant likelihood metrics and 5,420,745 SNPs of these SNPs exhibited minor allele frequencies >5%. Using the ~19 million quality-filtered SNPs, snpEff estimated the overall transition-transversion ratio at 1.89 (S4 Fig). Approximately 50% of the predicted variant effects fell into intergenic regions, with 20% occurring in upstream regions, 19% in downstream regions, and the remaining genic regions constituting 11%. Since most of the variant effects fell within intergenic regions, the variants that had low (451,663), moderate (439,159), and high impacts (15,019) accounted for a small proportion of the total variants. In total, 7,399 genes contained high-impact variants while low- and moderate-impact variants were observed across many annotated genes and accounted for 31,850 and 32,215 genes, respectively. To compare the coverage quality of our WGS data to existing GBS marker data, we binned variants in 5 kb windows across the genome. In our comparison, we observed that the GBS data did not have any variants for half of the bins across the sorghum genome whereas WGS had at least one variant in those bins (Fig 1A). Consistent with the methylation-sensitive nature of the ApeKI enzyme used to generate the GBS marker data, the GBS markers exhibited a strong bias towards genic regions while the distribution of variants in WGS data largely mirrored the overall proportion of genic and intergenic sequences in the sorghum genome with ~50% of total markers located in the intergenic regions, which were defined as the regions between annotated gene models or between a gene and the end of a chromosome (Fig 1B). In general, the WGS data showed higher variant density on chromosome arms and telomeric regions than in pericentromeric and centromeric regions (Fig 2A). The genome-wide average nucleotide diversity was 2.4 x 10-3 for the entire population with variation in SNP density across the genome showing telomeric regions accumulate more mutations than centromeric regions because of higher recombination rates (Fig 2A).

**Figure 1.**
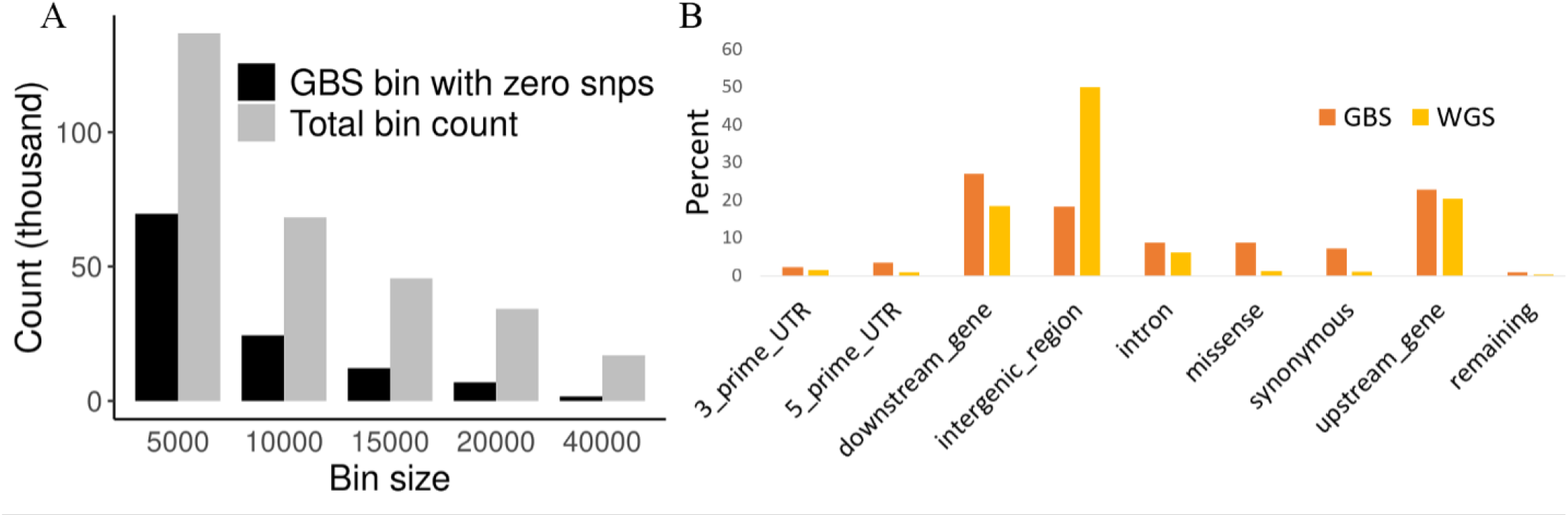
Comparison of Genotyping-by-Sequencing (GBS) and whole-genome sequencing (WGS) single-nucleotide polymorphism (SNP) distributions. Total bin count and counts for bins where GBS data lacked a SNP but WGS had SNPs across different bin sizes are shown in **A**, and **B** includes the percentage of variants across the major genic and intergenic regions for both GBS and WGS data.

**Figure 2.**
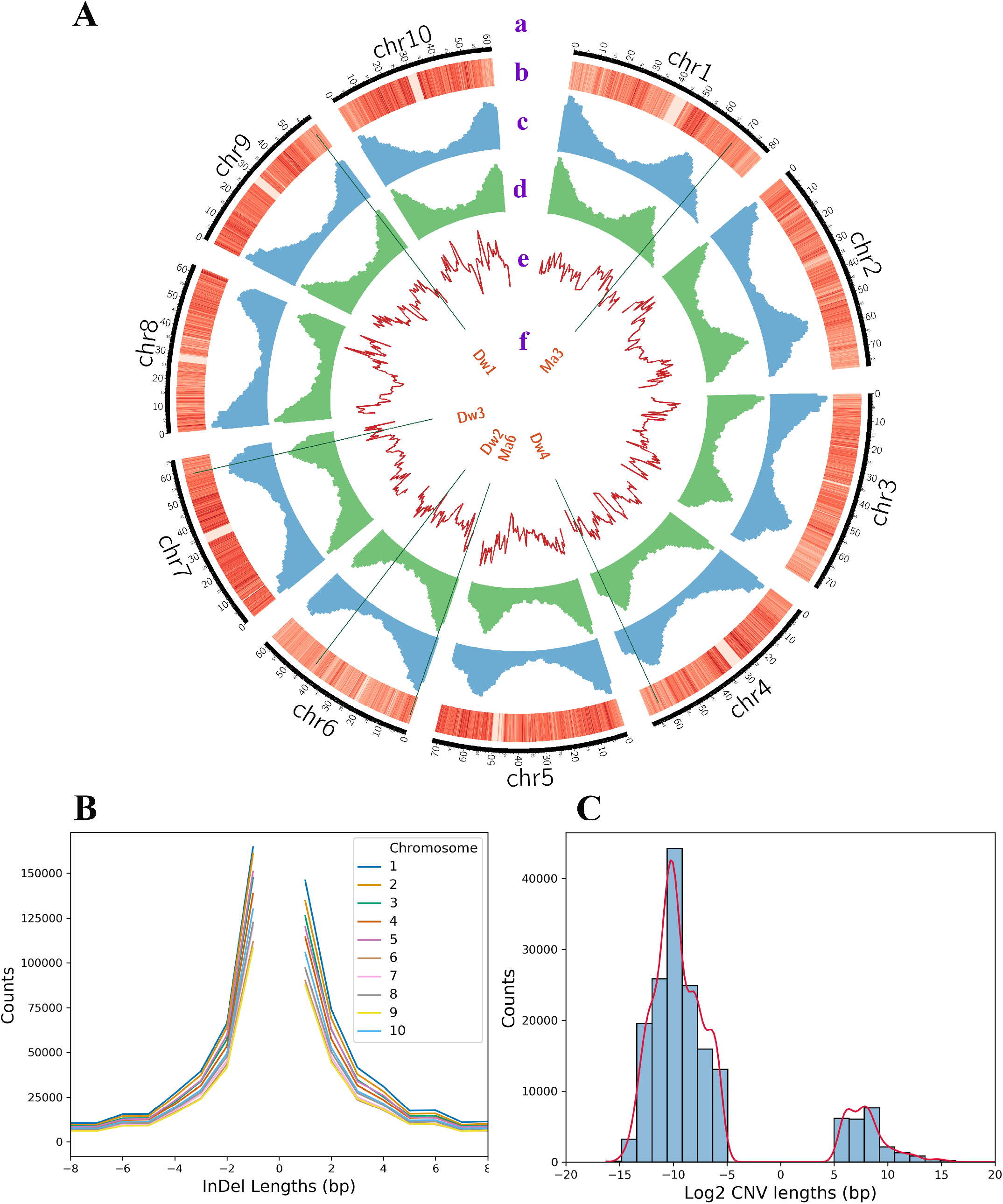
Genome-wide variant coverage and diversity. (A) Circos plot containing the sorghum karyotype, SNP density heatmap, indel histogram, copy number variant (CNV) histogram, nucleotide diversity, and genes in tracks a-f, respectively. (B) Line plot demonstrating total number of indels across varying indel lengths. (C) Histogram and kernel density estimate for CNV counts across varying log2 CNV lengths.

A total of 2,652,314 indels were identified after quality filtering with a strong telomeric bias in distribution (Fig 2A). The majority (82.56%) of indels were less than 15 bp in length (Fig 2B; S5 Fig), whereas the largest indel was 387 bp. Larger CNVs identified using Hecaton, ranging from 49 bp to 1 Mb, were retained for analyses, whereas all CNVs over 1 Mb in length were deemed false positives due to the limitations of short-read data to capture exceptionally large indels [45]. We identified nearly 7× more CNVs compared to a previous study in sorghum [61] likely due to the variant callers used as [61] only used LUMPY, but we used LUMPY and three other variant callers within Hecaton. The distribution of CNV types showed a preponderance of deletions over insertions (Fig 2C) and higher density around telomeric regions (Fig 2A), which is consistent with previous observation in sorghum [61].

### Population structure, haplotype blocks, and variant graph construction

We estimated the linkage disequilibrium (LD) decay distance for individual chromosomes as well as for the whole genome because LD influences genetic mapping resolution and is essential in haplotype construction. The genome-wide average distance at which LD decayed to *r*^2^ < 0.2 was approximately 20 kb, and the LD decay values plateaued around 150 kb (S6 Fig). Chr6 exhibited consistently higher LD compared to the other chromosomes, which is consistent with previous reports of limited recombination in Chr6 [62, 63] and as a result, the average physical size of estimated haplotype blocks was larger for Chr6.

Discriminant analysis (DAPC) estimated the optimal number of clusters to be eight, but there was no significant difference in Bayesian information criterion (BIC) for cluster counts between 6-11 (S7 Fig, S8 Fig). For simplicity, we opted to use the lowest number of population clusters (K=6) with lower BIC for subsequent ADMIXTURE analysis. The subpopulation grouping in the population structure analysis led to four clusters that correspond to the four botanical races of sorghum (caudatum, kafir, guinea, and durra) (S9 Fig). The fifth subpopulation cluster consisted of several durra-bicolor accessions that were historically categorized as milo and therefore we referred to that subpopulation/racial type as milo (S9 Fig). The sixth subpopulation consisted of some durra accessions but were mostly composed of mixed-race accessions and the accessions classified as bicolor, which is thought to be the early sorghum domesticate and therefore does not form a separate subpopulation cluster [14, 26]. The first 10 components in the principal component (PC) analysis accounted for about 36% of the genomic variation with the first three PCs explaining 9.36%, 7.86%, and 3.78% of the variation, respectively. The first PC separated kafir accessions from caudatum, PC2 separated kafir, caudatum and durra from the milo subgroup, and PC3 distinguished guinea accessions from all other accessions (Fig 3A,B). Ancestral population admixture was consistent with observed historical patterns among the sorghum races with greater admixture among approximately one fourth of the accessions (Fig 3C).

**Figure 3.**
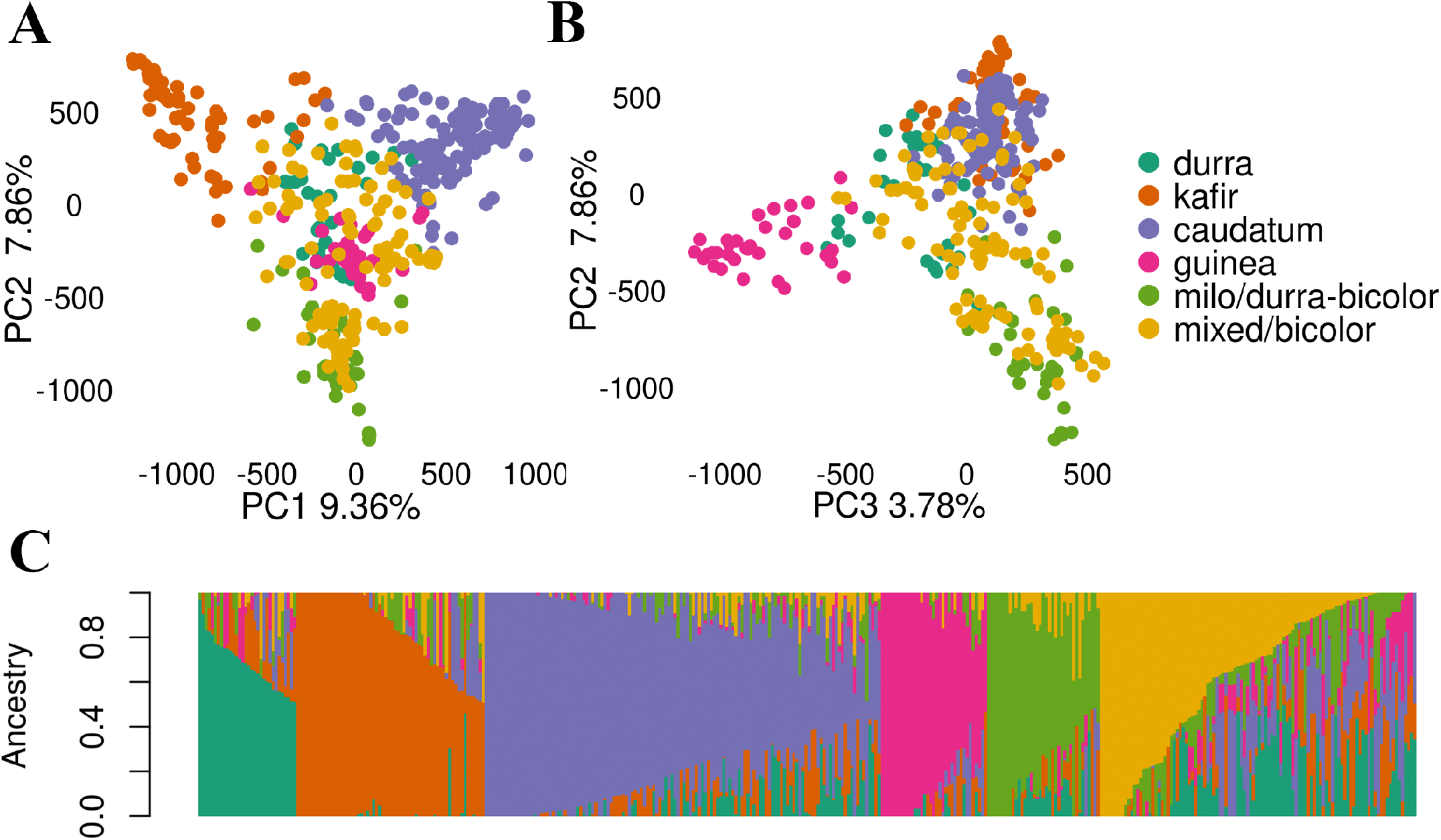
Population structure within the sorghum association panel using principal component analysis and admixture model (K=6). Subpopulations were labeled with corresponding botanical races or sorghum types that predominated for a given subpopulation.

A total of 35,029 haplotype blocks with an average length of 40 Kb were identified using Haploblocker[41]. Over 4,000 blocks were identified in chromosomes 2, 4, and 5, but fewer than 3,000 blocks in chromosomes 6, 7, and 9. Using the variant graph, we were able to successfully examine locus haplotype structure, map WGS reads from [60], and call SNPs, indels, and CNVs from the sample alignments. We obtained ~3 million variants per sample that subsequently reduced to ~1 million per sample following quality filtering. Compared to joint calling via the GATK pipeline and CNV calling using Hecaton, *vg* was able to call all variant types in a single run with a significantly shorter runtime than the alternative approaches employed above. However, the total number of variants obtained was an order of magnitude lower. This is likely due to several factors including increased power to detect variants when sharing haplotype and coverage information in joint calling [23], the use of multiple CNV callers by Hecaton [45], and the state of development for the variant calling methodology in *vg* [17].

### Genomic signatures of selection

The development of sorghum racial types is thought to be an outcome of multiple domestication events and subsequent local adaptations leading to the distinct population structure that is evident in our diversity panel as well [14, 64, 5]. Here, we wanted to identify and contrast signatures of historic selections during domestication to those that have occurred recently due to the photoperiod conversion of tropical accessions and/or selections made by breeding programs. We computed genome-wide F_st_ between racial subgroups arising from evolutionary diversification to identify signatures of historic selection, whereas genome wide F_st_ peaks between the latter groups (Converted and Bred) were used to identify signatures of artificial selection due to temperate conversion and/or breeding.

Several regions across the genome showed selective sweeps (F_st_ peaks) for subpopulations identified using population structure analysis (S10 Fig). There were 18 genomic regions with strong selection peaks, of which four regions across three chromosomes (Chr2, Chr3, and Chr8) had common F_st_ peaks in at least three subpopulations (Table 1, S10 Fig). The selected region around 45-54 Mb of Chr2 that had strong peaks for all but the kafir subpopulation have around 279 genes and seven quantitative trait loci (QTL) previously mapped to this region (Table 1). Among the seven QTLs in this region were three mapped for tannin by [65] and one each for amylose [66], panicle length [5], seedling survival [67], and anthracnose resistance [68]. Among the genes within this region, 24 genes had coiled-coil domains and showed significant enrichment based on the gene network analysis site, string-db. Another commonly selected region around 21-29 Mb of Chr3 had 68 genes that included several genes involved in biological regulation and molecular function including photosynthesis, but no QTL was previously mapped in the region (Table 1).

**Table 1.** Regions with strong (mean + 3 × standard deviation) selective sweeps based on F_st_ estimates for racial subpopulations identified from admixture analysis. Chr: Chromosome, QTLs: quantitative trait loci.

| Race/subpopulation | Chr | Start | End | Genes<sup>a</sup> | QTLs<sup>b</sup> |
| --- | --- | --- | --- | --- | --- |
| durra | 1 | 20.0 | 22.2 | 195 | 16 |
| mixed | 1 | 40.8 | 41.8 | 0 | 0 |
| milo | 1 | 70.3 | 71.3 | 193 | 43 |
| caudatum, durra, guinea, milo, mixed | 2 | 45.2 | 53.6 | 279 | 7 |
| guinea | 2 | 70.9 | 72.7 | 335 | 50 |
| milo | 2 | 76.9 | 78.7 | 159 | 0 |
| caudatum, durra, milo, mixed | 3 | 21.2 | 29.4 | 68 | 0 |
| durra | 3 | 14.1 | 15.2 | 74 | 26 |
| guinea | 3 | 17.1 | 19.6 | 93 | 0 |
| milo, guinea, caudatum | 3 | 40.2 | 45.7 | 88 | 0 |
| durra | 3 | 48.5 | 50.5 | 69 | 35 |
| caudatum | 4 | 64.4 | 66.0 | 255 | 71 |
| milo | 5 | 19.0 | 20.6 | 58 | 0 |
| kafir | 5 | 24.2 | 55.0 | 272 | 16 |
| caudatum, durra, milo | 8 | 7.8 | 11.4 | 121 | 14 |
| durra | 8 | 18.0 | 23.0 | 38 | 0 |
| guinea | 10 | 39.7 | 40.8 | 4 | 0 |
| milo | 10 | 61.1 | 62.2 | 19 | 2 |

<sup>a</sup> number of genes based on sorghum BTx623 v3.1.1 annotation (phytozome.org)
<sup>b</sup> number of QTLs within the genomic region based on Sorghum QTL atlas (aussorgm.org.au)

In general, the milo subpopulation had the highest number (8) of significant sweeps followed by durra (7) while kafir had the lowest (1) (Table 1). The only strong selection sweep identified in kafir was unique to the subpopulation and ranged from 24 to 55 Mb of Chr5 (Table 1). The caudatum subpopulation had a selection peak at 64-66 Mb of Chr4 near the *Tan1b* locus (Sobic.004G280800) that is associated with tannin/polyphenolic content [69]. Milo showed strong selection between 70-71 Mb of Chr1 which is near the Y locus (Sobic.001G397900) that encodes for yellow pericarp (seed color) in sorghum [70] and was captured using testa pigmentation (S10 Fig).

Durra, the subpopulation that is closely related to most of the accessions within the milo subpopulation, also had a minor peak around this region (S10 Fig). The only unique selection peak for the mixed subpopulation occurred around the non-genic region ranging from 41 to 42 Mb of Chr1. In general, most of these genomic regions with strong selection signatures had many characterized genes and several previously mapped quantitative trait loci (QTL) in and around these regions (Table 1).

Based on the F_st_ estimates, a total of six genomic regions showed strong selective sweeps between the accessions in the converted and bred groups (Fig 4A). The strongest peak was observed around 41 to 47 Mb of Chr6, this region contains major effect maturity (*Ma1* : Sobic.006G057866) and height (*Dw2* : Sobic.006G067700) genes that were introgressed for early maturity and short stature, respectively, during sorghum conversion (Fig 4A). Another region that showed a strong selective sweep was the region around the *Tan1* gene that is associated with tannin content (Fig 4A). Additional peaks were observed in the beginning of Chr1, Chr2, Chr4, and Chr8. The genomic region around the waxy locus (Sobic.010G022600) also showed a minor peak that was one standard deviation above the mean but did not reach two standard deviations above the mean (Fig 4A). We calculated expected heterozygosity (2pq) for converted and bred groups on a per-site basis using the allele frequencies (p and q) of SNPs at a given site. Figure 4B shows expected heterozygosity across each site for accessions in the converted group relative to the accessions in the bred category. The variation in relative heterozygosity for the two groups was consistent with the distribution of F_st_ peaks between the two groups (Fig 4A,B).

**Figure 4.**
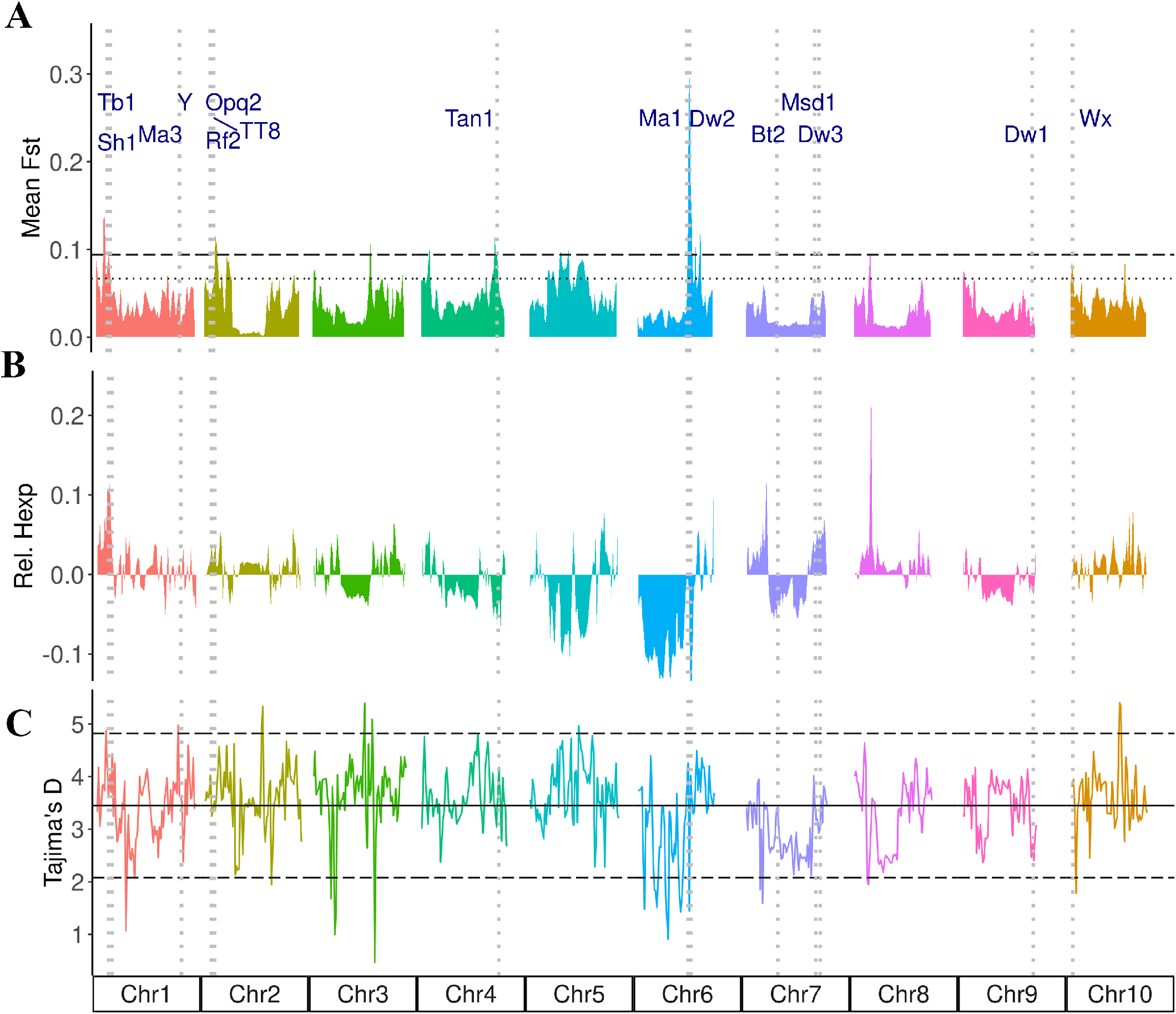
Genome-wide signatures of selection. **A** shows the mean F_st_ across the sorghum genome between tropical converted and temperate breeding acessions. **B** shows expected heterozygosity in converted group relative to the bred group. **C** shows the genome-wide Tajima’s D estimates. The horizontal lines show mean and standard deviations of the estimates: solid black lines show genome-wide average Tajima’s D, dotted lines show mean plus one standard deviation and long dashed lines show mean plus two standard deviation of the estimates. Vertical dotted lines show genes and loci related to height, maturity and other domestication related traits.

The genomic regions around the dwarfing and maturity genes showed strong bottlenecks for nucleotide diversity and Tajima’s D in the SAP (Fig 1A, 4C), and the whole-genome average value for Tajima’s D was 3.45, indicating that there are fewer rare alleles across the genome because of extensive inbreeding across the population. Most of the genomic regions showed Tajima’s D above the mean value indicating balancing selection while some regions, particularly at Chr1, Chr3, and Chr6, showed strong bottlenecks, indicative of purifying selection (Fig 4C). The regions in the middle of Chr6 and Chr7 showed a strong bottleneck in Tajima’s D and expected heterozygosity for converted lines compared to the breeding lines (Fig 4B, S11 Fig).

### Genome-wide association for plant height and tannin content

In sorghum, plant height (PH) and tannin content have been thoroughly examined due to their significant impacts on both historic selection [71] and modern agriculture [72]. Plant height in sorghum is genetically controlled by multiple genes including three predominant loci (*Dw1* : Sobic.009G229800, *Dw2* : Sobic.006G067700, and *Dw3* : Sobic.007G163800), which explain a large majority of the phenotypic variation in our population. All three major dwarfing loci showed significant association for PH using all three variant types independently (Fig 5A). For both SNP and indel variants, we also identified significant association for the previously reported *Dw4* locus, which occurs ~6.6 Mb on Chr6 [5, 73]. Additionally, we identified a significantly associated indel at ~6.5 Mb on Chr4 that overlaps with a previously identified QTL for total PH, flag-leaf height, and flag-leaf-to-apex interval (Fig 5A, Table 2) [74]. This locus contained eight genes; two of which were functionally annotated (Table 2). Among them, one gene was associated with plant viral response, and the other gene encoded an F-box protein, which is known to regulate plant vegetative and reproductive growth. A novel locus that is ~2 Mb downstream of maturity gene Ma3 (Sobic.001G394400) also showed a significant association for PH for SNP as well as indel variants (Fig 5A, Table 2). This locus is a hotspot for heat shock protein 70 (HSP70) with five HSP70 proteins occurring within 20 Kb and more than 10 HSP70 within 100 Kb of the associated SNP peak.

**Figure 5.**
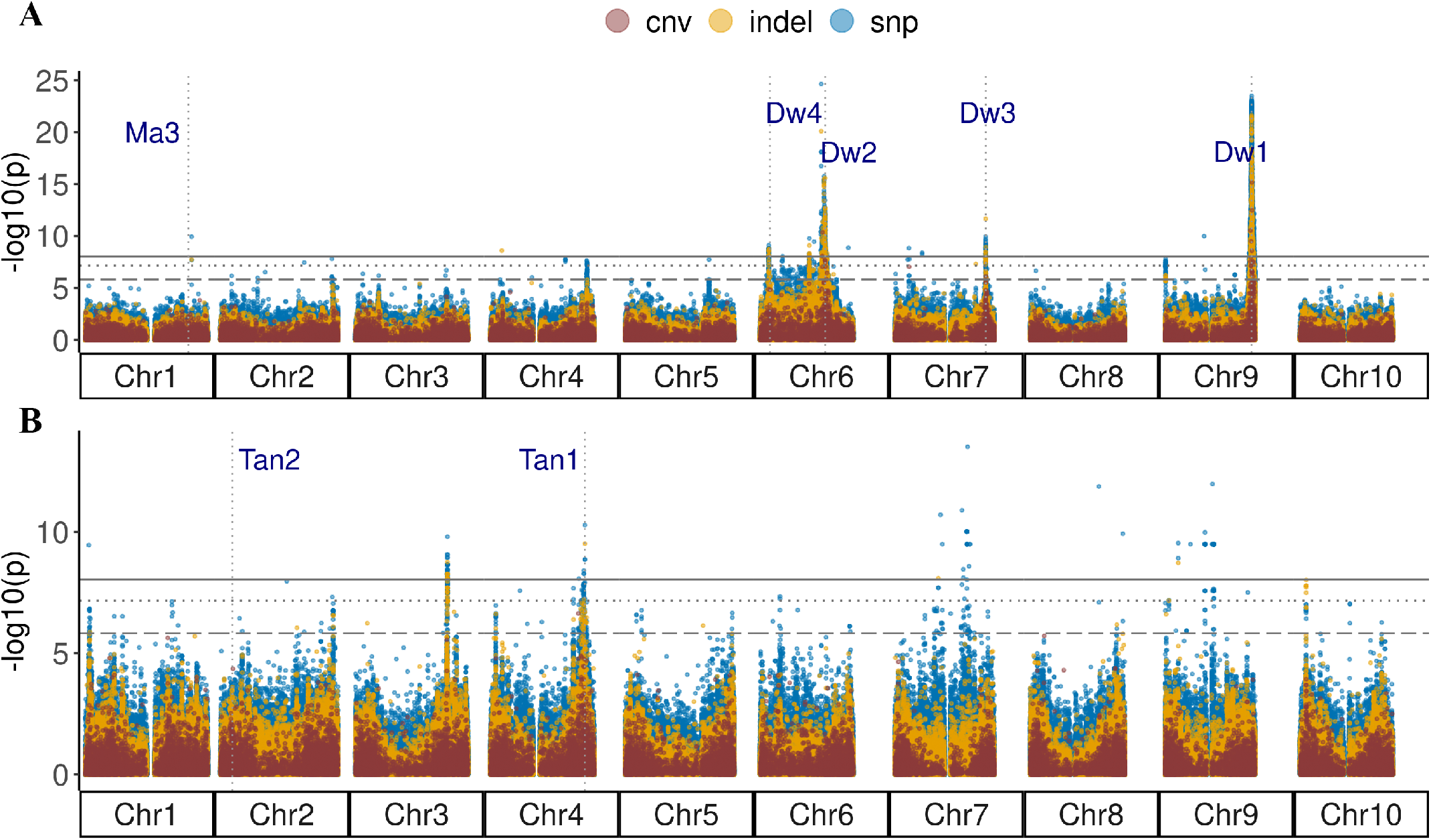
Genome-wide associations for plant height (A) and tannin content (B) using linear mixed models in GEMMA. Horizontal lines with solid, dotted, and dashed patterns represent Bonferroni-adjusted threshold of 0.05 for SNPs, indels, and CNVs, respectively. Vertical dotted lines indicate the positions of known genes and loci for height (Dw), maturity(Ma), and tannin (Tan).

**Table 2.** Putatively novel associated loci identified using whole-genome sequencing (WGS) data. Chr: chromosome, SNPs: single nucleotide polymorphisms, CNV: copy number variants, Chr: chromosome, p: p-values.

| Trait | Variant Type | Chr | Peak SNP | $-\log_{10}(p)$ | Gene Count<sup>a</sup> |
| --- | --- | --- | --- | --- | --- |
| Plant height | SNP | 1 | 69,987,248 | 9.93 | 4 |
| Plant height | Indel | 4 | 7,622,167 | 8.60 | 8 |
| Plant height | CNV | 7 | 9,225,709 | 7.04 | 1 |
| Tannin | SNP | 3 | 60,368,179 | 8.27 | 15 |
| Tannin | SNP | 3 | 60,722,769 | 9.80 | 6 |
| Tannin | SNP | 8 | 45,442,123 | 11.87 | 2 |
| Tannin | SNP | 8 | 61,186,817 | 9.92 | 9 |

<sup>a</sup> total genes within 20 kb of the peak SNP for the associated locus.

Tannin content is another important domestication trait. While higher tannin content lowers nutrient uptake [75], the presence of such phenolic compounds can conversely be important in reducing pest damage [71] as well as providing antimicrobial [76] and antioxidant activities that improve gut health [75]. One of the established primary regulators of tannin content is *Tannin1* (*Tan1*), which was identified in our GWAS for tannin content using all variant types (Fig 5B). Another important locus, *Tan2* (*TT8* : Sobic.002G076600), was not identified in GWAS using tannin content likely due to duplicate recessive epistatic interactions between the *Tan1* and *Tan2* loci [71]. However, when we conducted GWAS using phenotypic data indicating presence or absence of testa layer in our SAP accessions using a probit Bayesian sparse linear mixed model (BSLMM), the *Tan2* locus showed significant association using the SNP markers (S12 Fig). Two novel associations with strong peaks were identified for tannin content with SNPs, indels, and CNVs between 60 and 61 Mb of Chr3 (Fig 5B; S13 Fig). Previously, a significant association had been identified with 3-deoxyanthocyanidins around 59.7 Mb of Chr3 [65]. The novel loci at 60-61 Mb of Chr3 consist of several potential candidate genes that are involved in membrane transport, aromatic amino acid synthesis, and terpenoid pathways (Table 2). The top SNP at ~60.7 Mb was located within Sobic.003G270500, a gene encoding a farnesyl diphosphate transferase, which functions in the biosynthesis of terpenes and terpenoids [77]. Similarly, peaks on Chr7 at ~61.1 Mb and Chr9 at ~53.8 Mb were previously associated with proanthocyanidins [65], and the peak on Chr8 at ~61.1 has been identified with both polyphenol content and grain color [65].

### Pleiotropy analysis for grain yield and quality traits

Using the 25 traits collected by [15, 24, 25, 26], we first performed GWAS for all traits using LMMs in GEMMA [51]. From those initial LMM results, 19 traits were subsequently analyzed for pleiotropic effects using MashR [56]. MashR uses empirical Bayes methods to estimate patterns of similarity among conditions, and the resulting patterns are then used to improve the accuracy of effect estimates. Over 10,000 markers exhibited significant pleiotropic effects – nearly 16x more than was identified for over 100 traits using GBS data – across the sorghum genome with many well-known loci such as *Dw1*, *Dw2*, *Dw3*, *Ma1*, and *Ma3* (S14 Fig) exhibiting strong pleiotropic effects across multiple traits [78]. Many markers demonstrated an effect across 10 or more traits, and only Chr10 did not exhibit significant pleiotropic effects across more than five traits. Association results for various traits showed strong correlation between each other in pleiotropic analyses of grain yield and quality traits (S15 Fig, S16 Fig).

### Genome-wide prediction using WGS and GBS markers

Prediction results using WGS SNP data showed a significantly higher predictive ability (p-value < 2e-16) for all traits compared to GBLUP models using GBS SNP data (Fig 6). Predictive ability was, on average, 29% higher for WGS data and ranged from 13 to 47% across the traits studied. Mean predictive abilities ranged from 0.34 to 0.57 for GBS data and 0.44 to 0.71 for WGS data with thousand-grain weight and protein having the highest and lowest predictive abilities across both sequencing types, respectively. Among the traits, starch showed the largest (47%) increase in mean predictive ability from GBS to WGS, while days to anthesis showed the smallest (13%) increase.

**Figure 6.**
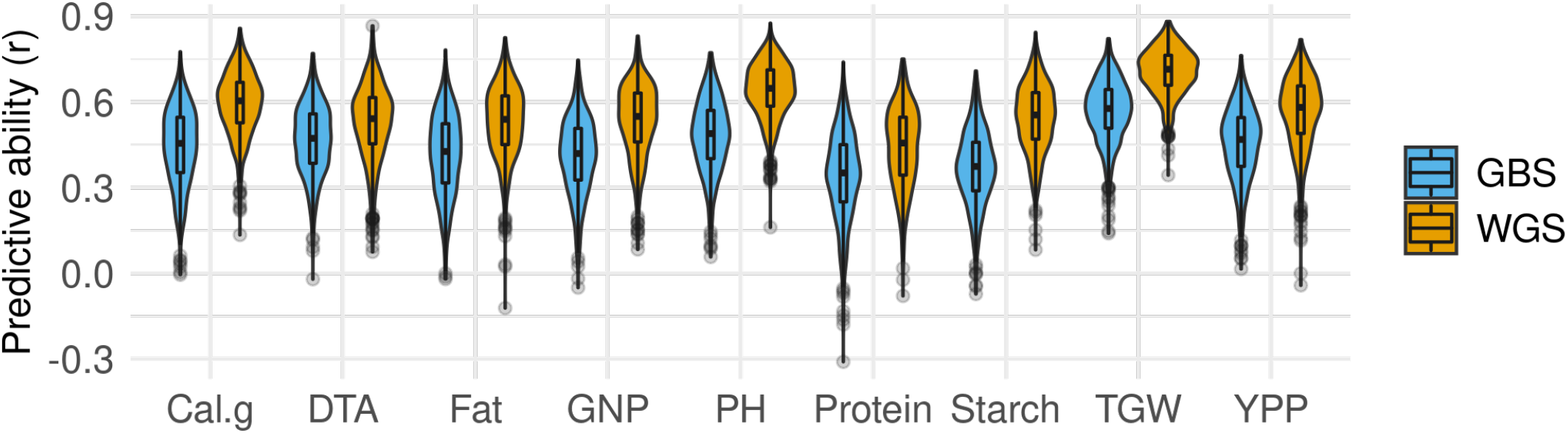
Predictive ability for genomic prediction of traits using SNPs from whole-genome sequencing (WGS) and genotyping-by-sequencing (GBS). Cal.g: Calories per gram, DTA: days to anthesis, GNP: grain number per primary panicle, PH: plant height, TGW: thousand grain weight, and YPP: yield per primary panicle.

## Discussion

Since its development, the United States Sorghum Association Panel has served as a pivotal resource for genetic dissection and as a source of genetic diversity for breeding [4, 79, 80]. To expand the breadth and depth of genomic data available for this crucial diversity panel, we sequenced the accessions in this panel at a higher depth and provide reliable high-density genome-wide markers for elucidating the genetic architecture of traits and propelling genomics-assisted breeding. Previously, this panel was characterized with GBS, which can be biased toward genic sequences, and therefore may misrepresent diversity across the population or in individuals within the population [81]. Here, with corroborating evidence, we demonstrate the inherent value of WGS data as demonstrated by reduced type-I and type-II errors, improved mapping resolution by capturing more recombination events, increased the depth of variants called, and the added benefits of different identifiable variants, which lead to improvements in genetic dissection and genome-wide prediction.

### High-density variants for genomic research and breeding

Apart from SNP markers, we identified a substantial number of insertions, deletions, and CNVs that contribute to our understanding of the complexity of the sorghum genome and the evolutionary processes that result in or develop from that complexity [82]. To date, there are only a few large-scale studies that have evaluated the utility of high-throughput indel data for GWAS and GP in human cohorts, while no such study was found in plants [83, 84]. Detection of SNPs is significantly easier than indel and CNV identification due to sequencing and reference biases, library preparation requirements, and algorithmic artifacts [85, 84]. Indels represent the second-most-common type of genetic variant. Yet, their value for identifying genome-wide associations has been overlooked due to limitations in both production cost and scalability [83]. Few studies in sorghum have examined indels and CNVs at scale [86, 87, 61, 88, 89], and none of these demonstrated the comparative value of all three variant types for GWAS. In fact, comparisons across all three variant types are limited to human studies where funds and scalability are less limiting, but even human studies lack a comprehensive review of the topic [83, 84]. Identification of indels and CNVs requires unbiased high-throughput sequencing or long-read sequencing to confidently call variants [90], and as the SAP was sequenced at ~38X, this dataset is uniquely suited to obtain high-quality variants of all three types. The number of SNPs generated in our study is consistent with the sequencing depth and population scale differences as previously reported for WGS in sorghum [60].

### Population structure, haplotypes, and variant graphs

Population structure analyses conducted using SSR markers were confirmed by the development of restriction-site-associated DNA sequencing, such as GBS [91, 4, 92, 5]. However, the true value of GBS was realized in downstream applications that showed increased mapping resolution for genome-wide association studies due to increases in genome coverage and ease of genotyping compared to SSR markers [92]. Similarly, results for population structure and genetic diversity analysis based on WGS data are similar to results based on GBS based markers despite increased marker density, which is not surprising considering SSR markers accurately captured population structure despite having much lower coverage than GBS markers [93]. Consistent with previous characterizations for the SAP, our population structure analysis subdivided the population into approximately six groups, which is consistent with the four botanical racial types, a milo subpopulation that includes the durra-bicolor of historic importance in breeding, and one admixed group that includes some durra race accessions, mixed-race accessions, and bicolor accessions that are thought to be the early domesticate and do not form a separate cluster [14, 91, 62, 26]. The average LD decay distance was approximately 20 kb (*r*^2^ < 0.2) for the whole genome but varied across chromosomes [94]. Notably, Chr6 failed to reach background levels (*r*^2^ < 0.2), which is consistent with previous results that found limited recombination on Chr6 [62, 63].

To date, there have been three sorghum pan-genomes published [18, 88, 89]. These pangenome construction efforts in sorghum have either utilized a smaller population size (N=176) and lower coverage (~10X) [18, 89] or called variants using sequence differences across multiple references [88]. Here, we identified more variants using higher coverage across more individuals, which will act as a pivotal resource for future pan-genomics in sorghum. Pan-genomes have the potential to provide significantly more information concerning potential haplotype structure across diverse panels than traditional reference genomes and can reduce the effects of reference sequence bias on read mapping and subsequent variant calling [20].

While some pan-genomic tools utilize pairwise alignment of multiple reference genomes to generate a pan-genome [95], the iterative alignment of reference genomes can result in a biased pan-genome [17]. The broad-scale high-throughput sequencing of diverse accessions can be foundational for development of variant graphs, particularly when the degree of large structural variants in a population is low. While variant graphs and pan-genomics are the future of reference-based genomics, we identified more quality variants using the GATK and Hecaton than from utilizing a variant graph approach with *vg*. Thus, while construction of a variant graph has the potential to reduce reference bias, the gains in bias reduction should be measured against the potential variant coverage that established variant callers can provide. To fully exploit the benefits of variant graphs, variants should be called using pangenomes constructed from multiple references so that contrasting haplotypes within a population or species can be captured.

### Distinct signatures for historic selection versus recent selection

Since the accessions in the SAP include various botanical races arising from evolutionary divergence and local adaptation, the genetic differentiation between racial types is indicative of differences arising from historic selection during domestication. Additionally, the accessions can also be divided differently into two groups: one group of individuals in the SAP is the converted lines from the SCP that were introgressed with maturity and height loci for photoperiod conversion and short stature [10, 11], whereas the other group of individuals includes the cultivars that were not only temperately adapted but were bred and selected through multiple generations and, as a result, new recombination events have allowed potentially different allelic combinations across the genome [11, 26].

The genome-wide signatures of selection were distinct for historic selection during domestication and local adaptation compared to selection signatures resulting from recent selection activities during photoperiod conversion and/or breeding (Fig 4). Since the converted lines were distributed across all botanical races, there were no distinct F_st_ peaks around the three genomic regions in Chr6, Chr7, and Chr9 that harbor the introgression for dwarfing and maturity genes into exotic tropical lines by the SCP. Thurber et. al. [96] had previously shown that introgression during the conversion process did not have any bearing on population structure analysis of converted lines. Based on the F_st_ peaks observed between the converted accessions and temperately adapted breeding lines, the haplotypes for the introgressed region in Chr6 could be distinct to the converted lines, whereas the other two introgressed regions in Chr7 and Chr9 show little differentiation with the breeding lines. This might be due to an abundance of recessive alleles for *Dw1* and *Dw3*, while the dwarfing allele for *Dw2* locus is rare among the breeding lines.

The genomic region in Chr2 (45-54 Mb) that showed common F_st_ peaks across all races exhibits a combination of positive selection at the beginning of the region with some bottleneck toward the end. The region at 53-54 Mb shows a strong bottleneck below two standard deviations of the mean Tajima’s D estimate, and an associated peak for anthracnose resistance was previously identified around 53.79 Mb of Chr2. This region contains a large (32 kb) sorghum gene, Sobic.002G169633, that encodes a protein kinase with NB-ARC as well as LRR domains and is located approximately 130 kb upstream of the associated GWAS peak [68]. The presence of 24 candidate genes with coiled-coil domains suggests that this region might have several mutations that have been independently, positively selected for across sorghum races during local adaptation. This region of the genome needs to be studied further because it shows strong selection and may be important for genetic dissection and breeding for biotic resistance.

Another region that showed signs of a selective sweep between converted and bred lines as well as within the caudatum subpopulation was the region around the Tan1 locus. The tannin loci have historically been subjected to bidirectional selection because of varied local herbivore threats and human taste sensitivity, resulting in natural variation around these loci across sorghum germplasm [71]. Caudatum accessions make up 45% of the SAP accessions that were reported to have a pigmented testa layer, whereas 32% of the remaining accessions with pigmented testa were from the mixed subpopulation, which also has several accessions closely related to caudatum (S1 File). Also, 73% of accessions with pigmented testa were from the converted group, whereas only 12% of accessions in the bred group were pigmented. This difference could result in large differences in allele frequency between the two groups around this region (S1 File).

### WGS markers improved genome-wide association and prediction over GBS markers

In addition to the common height loci (*Dw1* -*Dw3*) and tannin content loci (*Tan1*) identified previously using GBS markers, we identified novel associations across two loci for plant height and four loci for tannin content that were not identified previously using GBS markers. Previous studies have demonstrated that WGS improves both the mapping resolution and ability to identify novel associations over marker data derived from GBS [97]. Apart from the three well-characterized height loci (*Dw1*, *Dw2*, and *Dw3*), we also detected significant association for the putative *Dw4* locus at ~6.6 Mb of Chr6 as previously reported [5, 73]. A novel height association detected in Chr1 was located within 1 Mb of a PH QTL previously reported [98]. While associations at major loci for height overlapped for different variant types, the significant associations for plant height on Chr4 and Chr7 that were unique to indel data show that indel variants can overcome limitations of SNP data in detecting potential false negative associations. The tannin loci (*Tan1*) we have identified is consistent with previous association results in the SAP [65]. While the *Tan2* locus was undetected in our tannin content association like previous association analysis, we show that probit GWAS using a Bayesian sparse linear model for presence or absence of testa layer can detect both *Tan1* and *Tan2* loci. As both plant height and tannin content represent important phenotypes in sorghum breeding, the consistency in association results compared to previous studies provides validity of this newly developed genomic resource while novel associations show incremental advantage in genetic dissection.

We performed genomic prediction (GBLUP) using a genomic relatedness matrix derived from GBS or WGS SNPs to compare their predictive ability for agronomic, yield, and quality traits. An average of 29% increase in predictive ability was observed across nine traits, which is a substantial increase for most of these traits as they are quantitative and complex traits. This improvement in predictive ability is due to the ability of WGS markers to capture the additive genomic relationship better via increased density and coverage of genetic markers, subsequently improving the total genetic variance explained by the model [99]. And since genetic gain is directly proportional to selection accuracy, such improvements in accuracy of prediction will have a cumulatively positive effect on the long-term genetic gain across sorghum breeding programs [100].

## Conclusion

Approximately 44 million variants of diverse types were called using WGS of the SAP, and these markers represents a major increase in the density and variant types available for future sorghum studies and open the opportunity for detailed variant graphs, improved genomic prediction, and detection of novel loci facilitating sorghum improvement. These data are provided as a community resource for the continued development of this multi-purpose, climate-resilient crop.

## Supporting information

Supporting information are available at Figshare at xx.

S1 File. Accessions in the sorghum association panel along with metadata for population clusters, origin, racial classification, and testa pigmentation.

S1 Table. GATK variants and corresponding types. S1 Fig. Histograms of read coverage per sample.

S2 Fig. Cumulative genome coverage across samples.

S3 Fig. Guanine-Cytosine content distribution across samples.

S4 Fig. Total count of nucleotide substitutions across sorghum chromosomes.

S5 Fig. Indel length distribution.

S6 Fig. Linkage disequilibrium decay across each sorghum chromosome and across the genome.

S7 Fig. Cumulative variance explained across the SAP principal components.

S8 Fig. Discriminant analysis of principal components across varying values of k (number of clusters).

S9 Fig. Heatmap showing genomic relatedness between individual accessions within the sorghum association panel.

S10 Fig. Regions across the sorghum genome demonstrating selective sweeps for various subpopulations based on ADMIXTURE analysis.

S11 Fig. Measures of Tajima’s D across the genome within the photoperiod converted lines (Conv), breeding lines (Bred), and the whole population.

S12 Fig. Genome-wide association for presence of testa layer using probit Bayesian sparse linear mixed model.

S13 Fig. Heatmap for linkage disequilibrium around association peak for tannin content in chromosome 3.

S14 Fig. Genome-wide measures for pleiotropic effects of associated regions for 19 traits.

S15 Fig. Trait correlation across the sorghum genome for the 19 traits in the pleiotropy analysis.

S16 Fig. Genome-wide associations for grain yield components (A) and grain composition (B) using linear mixed models.

S17 Fig. Variant graph of *Dw1* locus at different aspects demonstrating macro- and micro-variations in the locus structure.

## Acknowledgments

We would like to thank Lindsay Shields for her assistance with tissue collection and lyophilization. Computational analyses were performed on Clemson University’s Palmetto Cluster, and we thank the staff who assisted with cluster and software management.

## Data availability

Raw WGS data will be available at the European Nucleotide Archive. VCFs will be accessible at the European Variation Archive, and scripts will be available on GitHub (github.com/jlboat/SAP).

## Author contributions

JLB and SS conceptualized, developed, and implemented the study design, and wrote the manuscript. JLB performed sequence analyses, variant data generation, and association analyses. SS performed population genomic analyses and genomic prediction. HJ performed sequence quality control analysis. SK, JCS, RB, and ZB acquired funding, provided supervision, and managed the project. All authors reviewed the manuscript and approved the manuscript for publication.

## Funding

This project was funded in part by the Department of Energy’s Advanced Research Project Agency award number DE-AR0001134 and the Department of Energy’s Office of Science award number DE-SC0020355. Any opinions, findings, conclusions, or recommendations expressed in this publication are those of the authors and do not necessarily reflect the views of the U.S. Department of Energy.

## Conflict of interest

JCS has equity interests in Data2Bio LLC, a company that provides genotyping services using sequencing technology. The authors declare no other conflicts of interest.

